# Molecular Docking Studies of Phytochemicals of *Allophylus serratus* Against Cyclooxygenase-2 Enzyme

**DOI:** 10.1101/866152

**Authors:** Kero Jemal

## Abstract

*Allophylus serratus* is a medicinal plant used traditionally as anti-inflammatory agent. The main objectives of this study are to identify phytochemical compounds that have anti-inflammatory properties from the leaf extracts of *Allophylus serratus* and to search for cyclooxygenase-2 (COX-2) enzyme inhibitors through molecular docking. From the GC-MS analysis of leaf extracts of this plant, various phytochemicals were identified. About 10of these phytochemical compounds were analyzed for their drug likeliness based on Lipinski’s rule of five and inhibitor property against the cyclooxygenase (COX-2) enzyme, a protein responsible for inflammation The phytochemical compounds which satisfy the Lipinski’s rule such as 1H-Benzocycloheptene, 2,4a,5,6,7,8-hexahydro-3,5,5,9-tetramethyl-,(R) and Sulfurous acid, dipentyl ester were subjected to docking experiments using AutoDock Vina. The results from molecular docking study revealed that 1H-Benzocycloheptene, 2,4a,5,6,7,8-hexahydro-3,5,5,9-tetramethyl-, (R)-, Sulfurous acid, dipentyl ester and 1,2-Benzenedicarboxylic acid, bis(2-methylpropyl) ester bind effectively to the active site region of COX-2 with a binding energy of −8.9, −8.4, and −7.9, respectively. The binding energy of the phyto-compounds were compared with the known antiinflammatory drug Diclofenac that inhibit COX-2 enzyme. It was found that the phytochemical compounds from leaf extracts of *Allophylus serratus* have strong inhibitory effect on COX-2 enzyme and as a result they have potential anti-inflammatory medicinal values. Thus the study puts forth experimental validation for traditional antidote and these phyto-compounds could be further promoted as potential lead molecule.

## 1. Introduction

Inflammation is our body’s natural reaction(an important physiological reaction) to injury caused by infectious agents, burn, toxic or physical, chemical or traumatic damages (Battu *et al*., 2011). It is a defense response of body characterized by pain, redness, heat, swelling, and loss of function(Riedel *et al*., 2014).Inflammation is an innate and adaptive immune systems normal body process that occur as a response to microbial pathogen infection, chemical irritation and tissue injury(Pan *et al*., 2010). The purpose of inflammation is to eliminate or limit the spread of injurious agent and repairing tissue (Nathan, 2002).The reaction comprises systemic and local responses (Mohamed *et al*., 2011). There are different components to an inflammatory response such as a complex array of enzyme activation, mediator release, cell migration, tissue breakdown and repair. These responses are aimed at host defense and usually activated in most disease condition.

The main mediators of inflammation and pain are proteins known as prostaglandins (Watson *et al*., 2000). The key enzyme which plays crucial role in the synthesis of prostaglandins is cyclooxygenase 2 (COX-2). Inflammatory process consists of the acute stage and chronic stage. The acute stage which occurs just after tissue damage is characterized by increase in blood vessels permeability, releaseof excessfluid, proteins and short period accumulation of white bloodcells (Westlund, 2006). In acute stage inflammation, the main mediators are histamine, serotonin, and cyclooxygenase 2 (COX-2)(Watson*et al*., 2000). If the acute inflammation is not controlled correctly the second chronic stage inflammation will occur. The chronicstage inflammation is mediated bymany inflammatory mediators which include Prostaglandin E_2_ (PGE2), nitric oxide(NO) and lipoxygenases. When the inflammation reacheschronic stage, it results in diseases such as peptic ulcers, systemic lupus, rheumatoid arthritis, asthma and cancer.

In tissues with no inflammation the level of Prostaglandin E_2_ (PGE2) is low. But during acute inflammation response, the level of PGE2 increases in the tissues with inflammation. Further increase in the level of PGE2 occurs as immune cells come to the tissues (Tilley *et al*., 2001). The enzyme responsible for the biosynthesis of PGE2 which induce inflammation and inflammatory response is cyclooxygenase 2 (COX-2) enzymes. Cyclooxygenase 2 (COX-2) stimulates biosynthesis PGE2 and increases its level in the tissue. COX-2 plays a vital role in conversion of arachidonic acid to prostaglandins (Ricciotti*et al*., 2011). The prostaglandins (PGE2) produced by the effect of cyclooxygenase 2 (COX-2) enzyme promote inflammation. The main target of anti inflammatory drugs is the enzyme COX-2. Inhibition of COX-2 can provide relief from the symptoms of inflammation and pain (Saqib, 2009).The most common non-steroidal anti-inflammatory drugs (NSAIDs) used in the treatment of inflammation are ibuprofen, naproxen, diclofenac, indomethacin, and ketoprofen(Warden, 2010). These drugs inhibit the expression of cyclooxygenase 2 (COX-2) enzymesresponsible for the biosynthesis of PGE2 (Vane and Botting, 1987). But, the long term use of these drugs have well known side effects on gastrointestinal tract(may cause gastric ulcers), cardiovascular system and kidney (Traversa*et al*., 1995). In addition, these drugs are very expensive. On the other hand many medicinal plants had been used since long time as anti-inflammatory without any side effects. At present, much attention has been given in the searching ofmedicinal plants with antiinflammatory activity which is not only without side effects but also cheap. These medicinal plants used as anti-inflammatory activity may lead to the discovery of new therapeutic agent that is not only used to suppress the inflammation but also used in diverse disease conditions where the inflammation response is amplifying the disease process.

Molecular docking is a method which predicts the preferred orientation of one molecule to a second when bound to each other to form a stable complex (Lengauer and Rarey, 1996).Knowledge of the preferred orientation is important to predict the strength of association or binding affinity between two molecules. The associations between biologically relevant molecules such as proteins, nucleic acids, carbohydrates, and lipids play a central role in signal transduction. Furthermore, the relative orientation of the two interacting partners may affect the type of signal produced (e.g., agonism versus antagonism). Molecular docking studies how to or more molecular structure fit together. Therefore, molecular docking is useful for predicting both the strength and type of signal produced. Characterization of the binding behavior plays an important role in rational design of drugs as well as to elucidate fundamental biochemical processes (Kitchen *et al*, 2004). The action of some harmful proteins produced in the body of humans could be prevented by finding an inhibitor which binds to that particular protein. Due to its ability to predict the binding-conformation of small molecule ligands to the appropriate target binding site Molecular docking is one of the most frequently used methods in structurebased drug design.

*Allophylusserratus* (Roxb.)Kurz, (Synonym *Allophyluscobbe*Raeuschel; *Allophylusedulis*Radlk) (Dharmani and Palit 2005), commonly known as Tippani in Hindi, belongs to the family Sapindaceae. It is a small tree or shrub found all over different parts of India. Traditionally this plant carries a strong ethno-pharmacological background and has been used as anti inflammatory, anti ulcer, totreatelephantiasis, oedma, and fracture of bones and gastrointestinal disorders such as diarrhea, anorexiaand dyspepsia(Umashanker *et al*., (2011); Gupta and Tandon, 2004); Dharmani *et al*., (2005) and Kumar *et al*., (2010), reported that the ethanolic extract of *Allophylus serratus* has potential anti ulcerogenic and anti osteoporotic activities respectively. The leaves are used to reduce fever, to relieve rashes, promote lactation, to treat colic to relieve stomach aches, as antiulcer and to reduce piles (Umashanker and Shruti, 2011; Devi *et al*., 2013). The roots of this plant contain tannin and are considered astringent and used for treating nose bleeding, diarrhea and rheumatic pains (Umashanker and Shruti, 2011).

The presences of different phytochemicals such as steroids, glycosides, flavonoids, alkaloids and phenolics in this species have been reported (Sanmuga *et al*., 2012). Phytochemical screening and Pharmacognostic studies of *Allophylus serratus* showed the presence of various chemical compounds in different parts of the plant. Leaves of the plant contain ß-sitosterol. They also contain phenacetamide, a chemical known for its antiulcer activity (Rastogi and Mehrotra, 1995). The presence of Quercetin, Pinitol, Luteolin-7-O-B-D-glucopyranoside, rutin, apigenin-4-O-B-D-glucosid also reported by (Kumar *et al*., 2010).

The aim of present study is to screen the phytochemicals present in the leaves extracts of *Allophylus serratus* by GC-MS analysisand to identify potential leading compounds of *Allophylus serratus* leaf extracts against Cyclooxygenase-2 protein target involved in inflammation by carrying out molecular docking studies.

## 2. Materials and Methods

### 2.1. Plant materials

Fresh, healthy and mature leaves of *Allophylus serratus* were collected from Andhra University, Visakhapatnam, India and authenticated by at Department of botany, Andhra University, Visakhapatnam. The voucher specimens (21921) were deposited in the herbarium, college of Science and Technology, Department of Botany, Andhra University.

### 2.2. Preparation of extract

The leaves of *Allophylus serratus were* thoroughly washed in tap water, dried under shade for one week and powderedin an electric mixer grinder to powder. 100 g of shade dried powder was extracted by soxhlet extraction with 500 mL of Methanol and Ethyl acetate separately. The solvents were then evaporated by using rotary evaporator and the extracts were concentrated to thick mixture. Crude extract obtained were keptat 4°C until further assay.

### 2.2. GC-MS(Gas chromatography-mass spectrometry)analysis of the extracts

GC-MS analysis was performed using SCHIMADZU(GCMS-QP2010 PLUS) andcarried out on a DB 5 – MS(0.25X30X0.25) capillary standard non - polar column and gas chromatograph interfaced to a mass spectrometer (GC-MS) instrument. The electron impact mode (electron ionization system with ionization energy)at 70 eV was operated and helium (99.999%) was used as carrier gas at a constant flow of 1 mL/min. The oven temperature was programmed from 45°C (isothermal for 4 min) with an increase of 10°C/min to 175°C, then 5°C/min to 240°C, ending with a 9 min isothermal at 240°C. Mass spectra were taken at 70eV; a scan interval of 0.5 seconds and fragments from 40–500Da. Total GC running time was 60 minutes. The relative percent amount of each component was calculated by comparing its average peak area to the total areas.

### 2.4. Identification of phytochemical Components

To interpret the mass spectrum of GC-MS, unknown components were compared with the spectrum of the known components using the databases of National Institute Standard and Technology version (NIST08s) WILEY8, FAME. The name, molecular weight and structure of the unknown components of the test materials were determined.

### 2.5. Molecular Docking Studies

Molecular docking is considered as the “key and lock” hypothesis used to find the best fit orientation of ligand and protein. Phytochemicals isolated from *Allophylus serratus* leaf extracts were selected for molecular docking study. Target protein Cyclooxygenase (COX-2) was docked with selected phytochemical compounds using UCSF Chimera (version 1.11.2) and AutoDock Vina software and binding energies were calculated. The ligands and the target protein were prepared following the standard procedure of ligand and protein preparation and the prepared files of the protein and ligands were submitted to AutoDock vina. The binding energy and the binding contacts of each ligand were obtained and the docked complexes were analyzed using Discovery Studio 3.1 visualizer.

#### 2.5.1. Ligand preparation

For the present study, from 42 phytochemical compounds or bioactive compounds identified from *Allophylus serratus* leaf extract by GC-MS analysis only ten of them were selected for the docking study. The chemical structures of all the phyto-compounds obtained from the result of GC-MS and selected for docking, were retrieved through the PubChem compound database at NCBI [http://pubchem.ncbi.nlm.nih.gov/].The 2D structures were obtained from PubChem compound database at NCBI and the 3D structure of the ligands was drawn by using UCSF Chimera. Ligands were prepared (minimization of energy done, hydrogen atoms added and charges added where required) using UCSF Chimera structure build module. The prepared 3D structures of the compounds were saved in the pdb format and were finally optimized for docking using UCSF Chimera tools.

#### 2.5.2. Retrieval of target protein

Cyclooxygenase-2 (COX-2) protein which plays a crucial role in modulation of inflammation was selected for its interaction with the phyto-constituents isolated from *Allophylus serratus* leaf extracts. The 3DX-ray crystal structure of the Cyclooxygenase-2 (COX-2)protein with details resolution was retrieved from Protein Data Bank (PDB) with PDB ID 6COX_A. The 6COX_A is a complex of COX-2 with an inhibitor SC-558 (http://www.rcsb.org/pdb).

#### 2.5.3. Preparation of the target protein

The raw PDB protein structure could not be used for molecular docking studies. PDB structure consists only of heavy atoms, waters, cofactors, metal ions and can be of multimeric. These structures do not have the information about bond orders, topologies or formal atomic charges. The terminal amide groups may be misaligned because the X-ray structure analysis cannot distinguish between O and NH2 Ionization and tautomeric states are also unassigned. So, the raw PDB structure retrieved from PDB should be prepared in a suitable manner for docking.

Before proceeding to docking analysis, theCOX-2 enzyme protein was subject to refinement and energy optimization. Protein Preparation Wizard of UCSF Chimera (Dockprep) software was used to process and prepare the protein. This Wizard allows one to properly prepare a protein for docking. This tool can convert a raw PDB structure into all-atom fully prepared protein models.

The X–ray crystal structure of 6COX_A protein was prepared by removing all the water molecules present in the structure. Since the raw data do not contain any hydrogen in it, the implicit hydrogen atoms were added to the atoms to satisfy their appropriate valences and ligands and ions of no significance present in the protein structure were deleted. Then the structure was optimized by assigning the bond orders, bond angles and topology. The formal atomic charges were fixed for the amino acid residues and energy minimization was carried out. The Discovery Studio 3.1 visualizer was used to analyze the protein structure, the hydrogen bond interactions and non bond interactions of ligands with the active site residues and preparation of high resolution images.

#### 2.5.4. Docking

After the ligands and the target protein prepared for docking, AutoDock Vina was used to perform docking process by bringing the ligand with the target protein. Individual ligand compound was given as input in the parameter meant for “ligand” and the protocol was run for each of the ligands. The various conformations for ligand in this docking procedure were generated and the final energy refinement of the ligand pose occurred The Dock score of the best poses docked into the target protein for all the tested compounds was calculated.

## 3. Results and Discussion

### 3.1. GC-MS Analysis

GC-MS analysis was performed for the methanolic and ethyl acetate leaf extracts of *Allophylus serratus* to assess their phytochemical constituents. The results of GC-MS analysis showed the identification of a number of phytochemical compounds. The compositions of the phytochemical compounds present in ethyl acetate and methanolic extracts of *Allophylus serratus* leaf extracts identified by GC-MS analysis with their retention time (RT), molecular formula, molecular weight and area (%) are presented in (Table 1). The GC-MS chromatograms of the two extracts are also given in Fig.1 and Fig 2 respectively.

**Figure 1:**
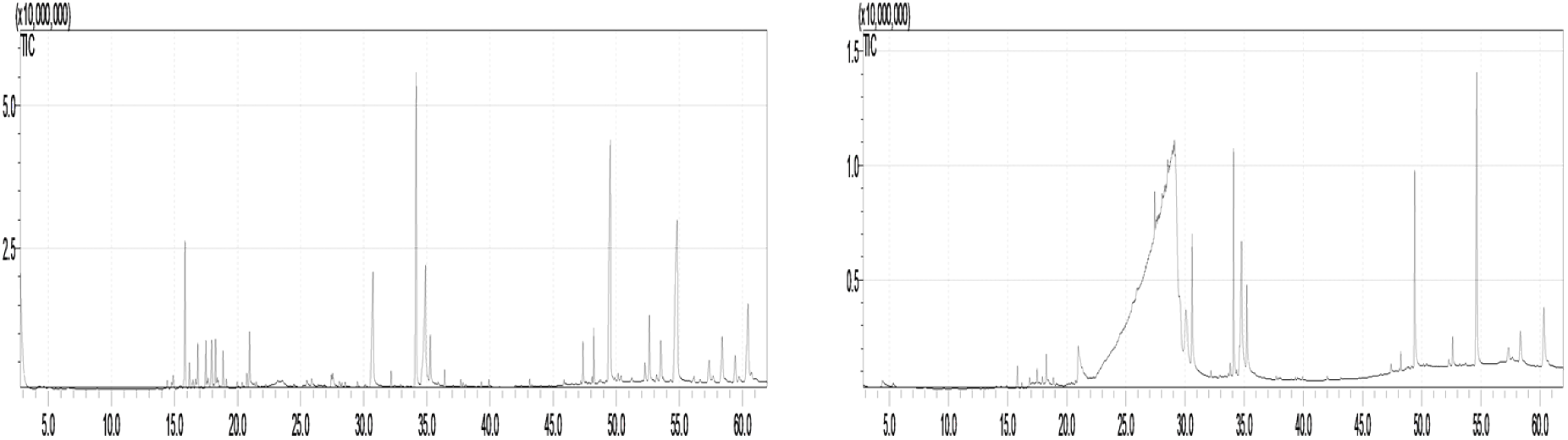
GC-MS chromatogram of methanol extract of *Allophylus serratus* leaves. **Figure 2:** GC-MS chromatogram of Ethyl Acetate extract of *Allophylus serratus* leaves

**Figure 2:**
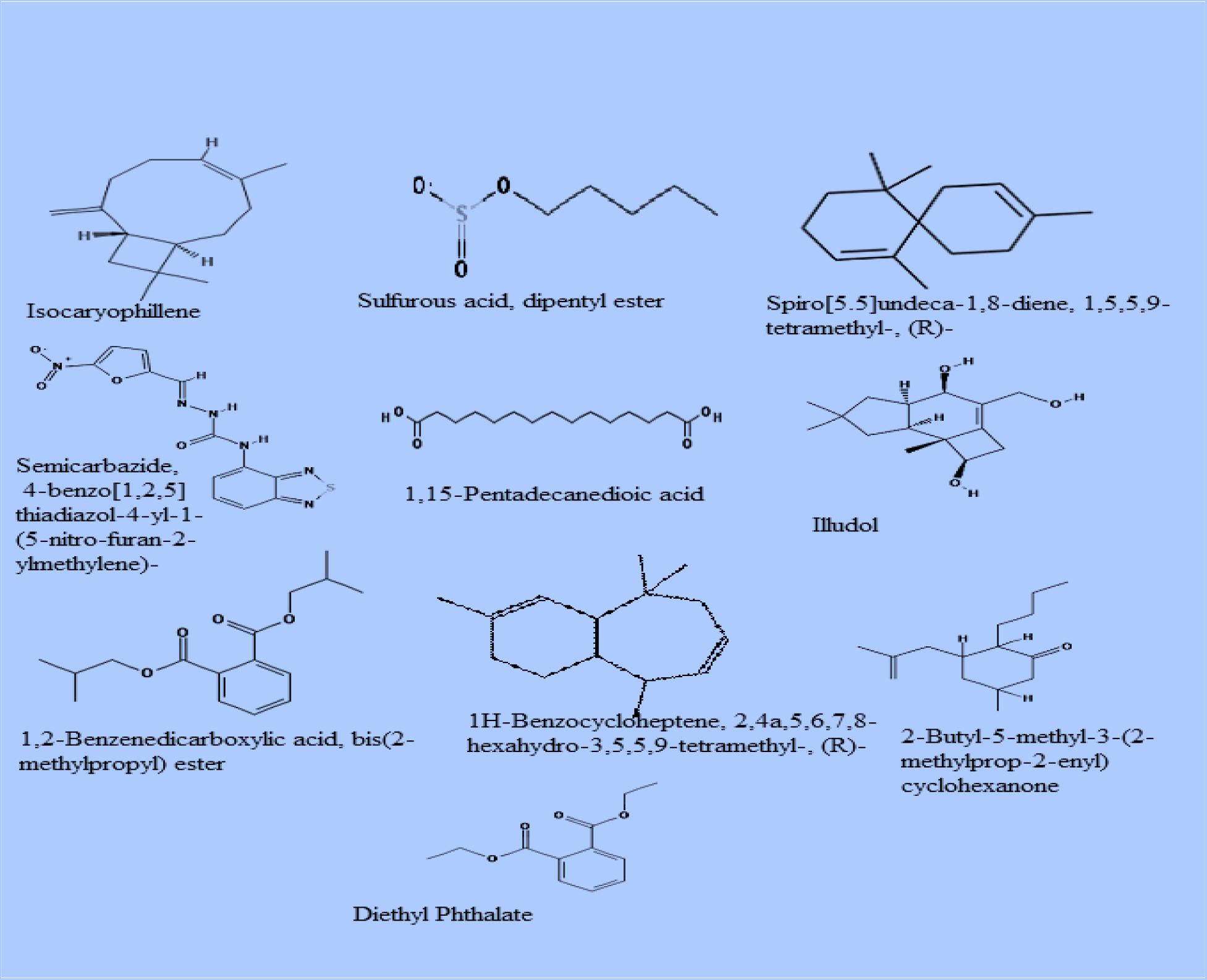
Structure of phytochemicals isolated from *Allophylus serratus* plant and used for molecular docking study.

**Table 1:**
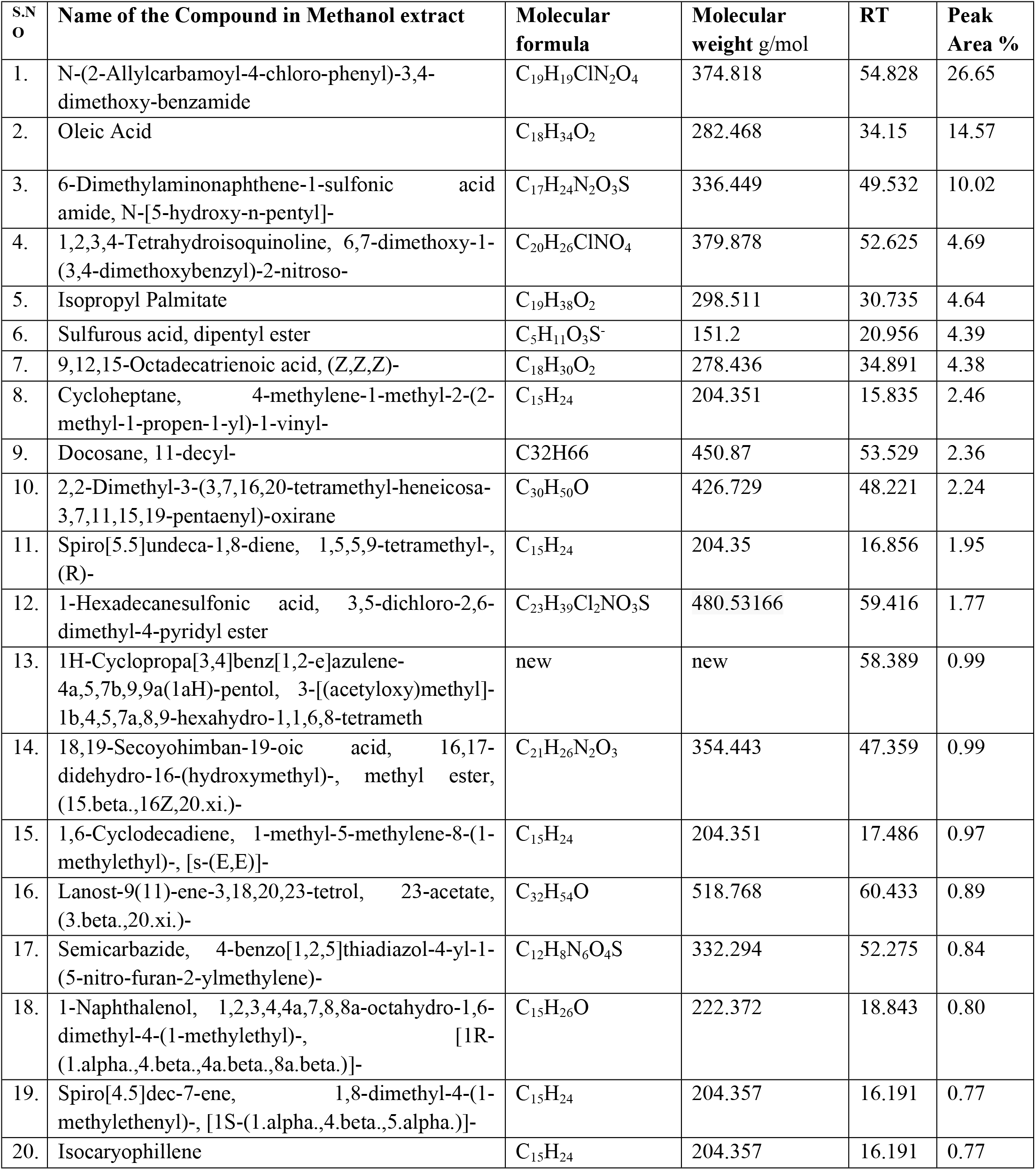
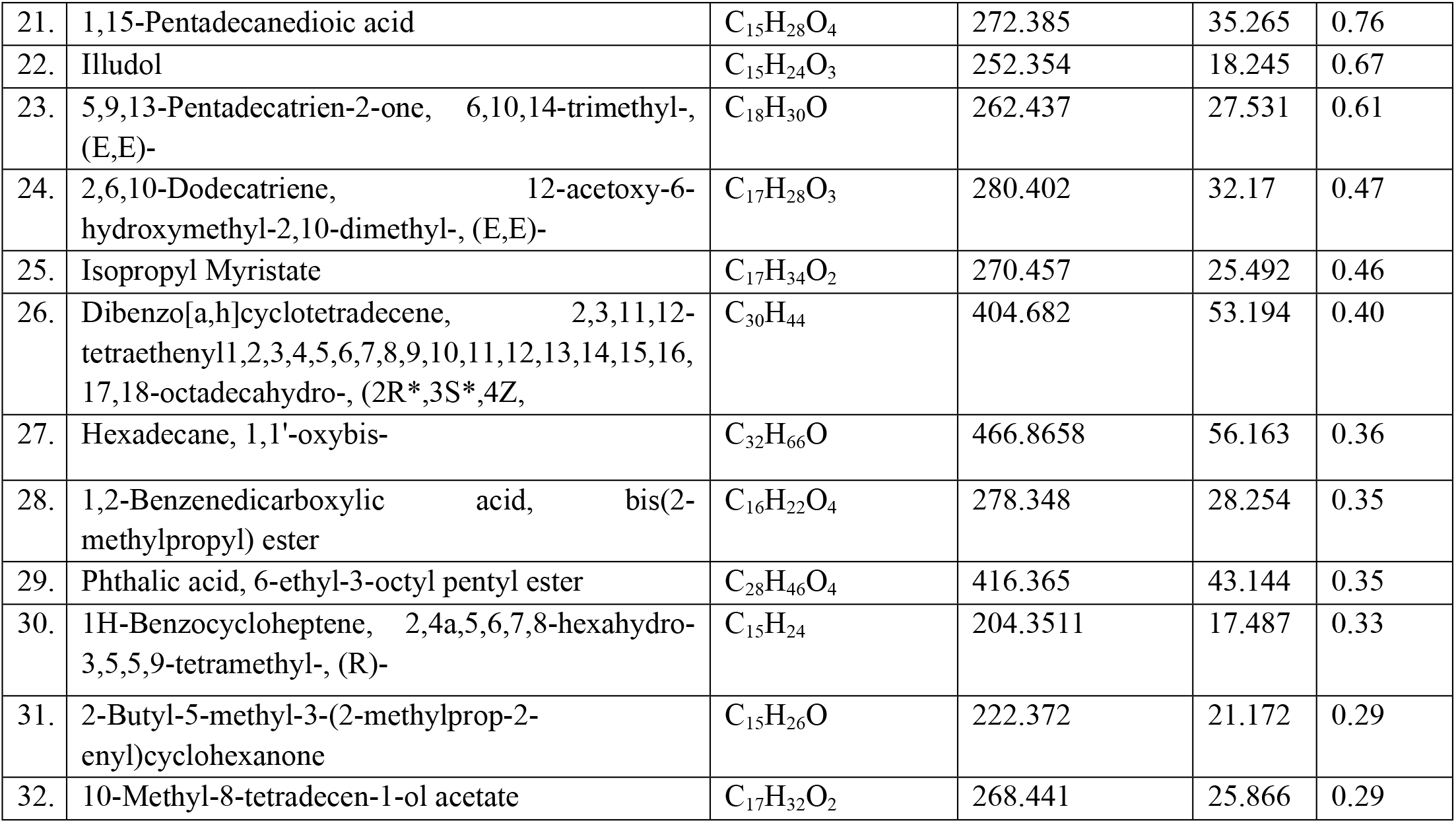
Bioactive compounds identified in the methanolic extract of leaf of *Allophylus serratus* by GC-MS analysis

GC–MS chromatogram of the methanol extract of leaves of *Allophylusserratus clearly* showed thirty two peaks indicating the presence of total thirty two compounds and other many compounds in smaller quantities (Fig.1a). Among the identified compounds in methanolic extract, the maximum percentage (26.65%) was that of N-(2-Allylcarbamoyl-4-chloro-phenyl)-3,4-dimethoxy-benzamide followed by Oleic Acid (14.57%),6-Dimethylaminonaphthene-1-sulfonic acid amide, N-[5-hydroxy-n-pentyl]- (10.01%), Isopropyl Palmitate (4.69%), Sulfurous acid, dipentyl ester (4.64%), 9,12,15-Octadecatrienoic acid, (Z,Z,Z)- (4.38%) (4.38%), Cycloheptane, 4-methylene-1-methyl-2-(2-methyl-1-propen-1-yl)-1-vinyl-(4.38%). The other compounds are found in small quantities as indicated in the Table 1. The bioactive compounds in the methanol leaf extracts identified by GC-MS with their Molecular formula, Molecular weight (MW), Retention time (RT), and Concentration (%)were presented in (Table 1).

GC–MS chromatogram of ethyl acetate extract showed twelve peaks indicating the presence of twelve major compounds (Fig1b and table1). Among the identified compounds in ethyl acetate extract, the maximum N2-Veratroylglycine N’-(4-fluoro-a-methylbenzylidene)hydrazide (27.36 %) followed by Sulfurous acid, 2-pentyl undecyl ester (10.63 %), Diethyl Phthalate (10.10 %), d-Glycero-d-tallo-heptose (9.67%), Galacto-heptulose (9.07%), 1,3-Dimethyl-5-[1,2-dicarbethoxyhydrazino]-6-hydrazinou%), 1,2,3,4-Tetrahydroisoquinoline, 6,7-dimethoxy-1-(3,4-dimethoxybenzyl)-2-nitroso-(2.66 %), Isopropyl racil (6.18 %), Isopropyl Palmitate (3.48 %), 9,12,15-Octadecatrienoic acid, (Z,Z,Z)- (3.35 stearate (2.05 %), 6-O-Methyl-2,4-methylene-.beta.-sedoheptitol (1.88 %) and 2,2-Dimethyl-3-(3,7,16,20-tetramethyl-heneicosa-3,7,11,15,19-pentaenyl)-oxirane (0.97 %). The bioactive compounds in the ethyl acetate leaf extracts identified by GC-MS with their Molecularformula, Molecular weight (MW), Retention time (RT), and Concentration (%)were presented in (Table 2).

**Table 2:**
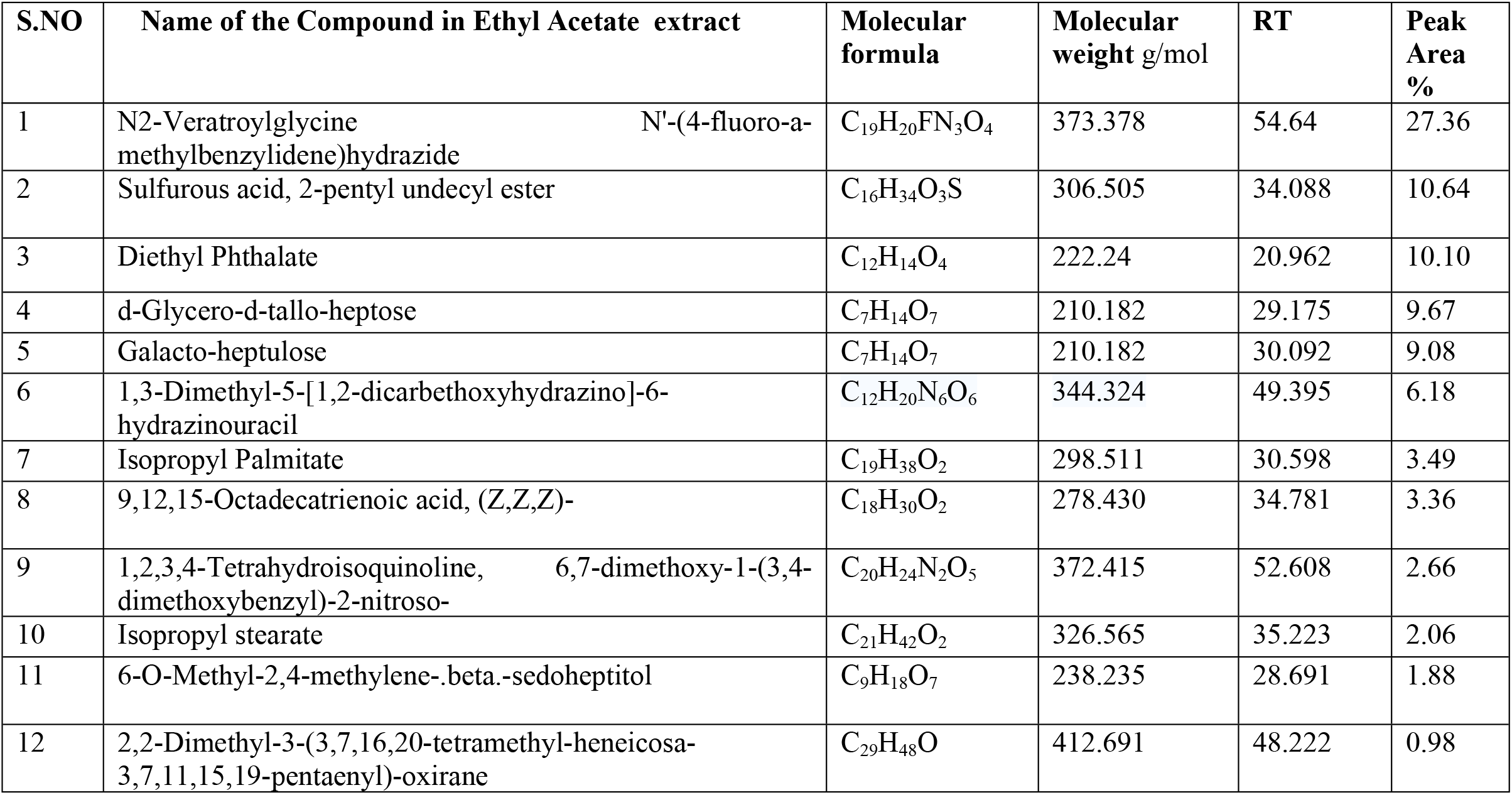
Bioactive compounds identified in the ethyl acetate extract of leaf of *AllophylusserratusbyGC-MS* analysis

| S.NO | Name of the Compound in Ethyl Acetate extract | Molecular formula | Molecular weight g/mol | RT | Peak Area % |
| --- | --- | --- | --- | --- | --- |
| 1 | N2-Veratroylglycine N'-(4-fluoro-a-methylbenzylidene)hydrazide | C<sub>19</sub>H<sub>20</sub>FN<sub>3</sub>O<sub>4</sub> | 373.378 | 54.64 | 27.36 |
| 2 | Sulfurous acid, 2-pentyl undecyl ester | C<sub>16</sub>H<sub>34</sub>O<sub>3</sub>S | 306.505 | 34.088 | 10.64 |
| 3 | Diethyl Phthalate | C<sub>12</sub>H<sub>14</sub>O<sub>4</sub> | 222.24 | 20.962 | 10.10 |
| 4 | d-Glycero-d-tallo-heptose | C<sub>7</sub>H<sub>14</sub>O<sub>7</sub> | 210.182 | 29.175 | 9.67 |
| 5 | Galacto-heptulose | C<sub>7</sub>H<sub>14</sub>O<sub>7</sub> | 210.182 | 30.092 | 9.08 |
| 6 | 1,3-Dimethyl-5-[1,2-dicarbethoxyhydrazino]-6-hydrazinouracil | C<sub>12</sub>H<sub>20</sub>N<sub>6</sub>O<sub>6</sub> | 344.324 | 49.395 | 6.18 |
| 7 | Isopropyl Palmitate | C<sub>19</sub>H<sub>38</sub>O<sub>2</sub> | 298.511 | 30.598 | 3.49 |
| 8 | 9,12,15-Octadecatrienoic acid, (Z,Z,Z)- | C<sub>18</sub>H<sub>30</sub>O<sub>2</sub> | 278.430 | 34.781 | 3.36 |
| 9 | 1,2,3,4-Tetrahydroisoquinoline, 6,7-dimethoxy-1-(3,4-dimethoxybenzyl)-2-nitroso- | C<sub>20</sub>H<sub>24</sub>N<sub>2</sub>O<sub>5</sub> | 372.415 | 52.608 | 2.66 |
| 10 | Isopropyl stearate | C<sub>21</sub>H<sub>42</sub>O<sub>2</sub> | 326.565 | 35.223 | 2.06 |
| 11 | 6-O-Methyl-2,4-methylene-.beta.-sedoheptitol | C<sub>9</sub>H<sub>18</sub>O<sub>7</sub> | 238.235 | 28.691 | 1.88 |
| 12 | 2,2-Dimethyl-3-(3,7,16,20-tetramethyl-heneicosa-3,7,11,15,19-pentaenyl)-oxirane | C<sub>29</sub>H<sub>48</sub>O | 412.691 | 48.222 | 0.98 |

### 3.2. Molecular Docking Results

#### 3.2.1. Ligand preparation

From the list of GC-MS identified chemical compounds in the methanol and ethyl acetate extracts of leaf of *Allophylus serratus* the following compounds (1) Isocaryophillene, (2) Sulfurous acid, dipentyl ester, (3) Spiro[5.5]undeca-1,8-diene, 1,5,5,9-tetramethyl-, (R)-, (4) Semicarbazide, 4-benzo[1,2,5]thiadiazol-4-yl-1-(5-nitro-furan-2-ylmethylene, (5)1,15-Pentadecanedioic acid, (6)Illudol, (7) 1,2-Benzenedicarboxylic acid, bis(2-methylpropyl) ester, (8) 1H-Benzocycloheptene, 2,4a,5,6,7,8-hexahydro-3,5,5,9-tetramethyl-, (R)-, (9) 2-Butyl-5-methyl-3-(2-methylprop-2-enyl)cyclohexanone and (10) Diethyl Phthalate were selected and subjected to docking experiments (Table 7.3). Diclofenac was used as standard control in the docking experiment. The 2D structure of and phytochemical properties of phytochemical compounds identified by GC-MS analysis in the leaf extracts of *Allophylus serratus* were downloaded from PubChem database (https://pubchem.ncbi.nlm.nih.gov/)(Table 3 and Fig 2). All the compounds were prepared for docking using the UCSF Chimera software version 1.11,2. During preparation of the ligand, 3D conformations were generated using UCSF hydrogen atoms were added and Gasteiger charges were added. The energy minimization was done running a 1000 cycles by steepest descent approximation and was converged to a gradient of 0.02 using the tool UCSF Chimera 1.11.2., and the AMBERff99SB Force field was used for this procedure. The ligands were saved as mol2 document and input ligand file format was mol2 for all docking programs investigated.

**Table 3:** Lipinski’s properties of the selected phytochemical compounds fromleaf extracts of*Allophylusserratus*

| SN | Compound Name | Molecular weight(<500KD) | Log P (<5) | H-bond donor (<5) | H-bond acceptor (<10) |
| --- | --- | --- | --- | --- | --- |
| 1 | Isocaryophyllene | 204.36 | 4.4 | 0 | 0 |
| 2 | Sulfurous acid, dipentyl ester | 151.20 | 1.1 | 0 | 4 |
| 3 | Spiro[5.5]undeca-1,8-diene, 1,5,5,9-tetramethyl-, (R)- | 204.35 | 4.4 | 0 | 0 |
| 4 | Semicarbazide, 4-benzo[1,2,5]thiadiazol-4-yl-1-(5-nitro-furan-2-ylmethylene)- | 332.29 | 2 | 2 | 8 |
| 5 | 1,15-Pentadecanedioic acid | 272.39 | 4.8 | 2 | 4 |
| 6 | Illudol | 252.35 | 1.2 | 3 | 3 |
| 7 | 1,2-Benzenedicarboxylic acid, bis(2-methylpropyl) ester | 278.35 | 4.1 | 0 | 4 |
| 8 | 1H-Benzocycloheptene, 2,4a,5,6,7,8-hexahydro-3,5,5,9-tetramethyl-, (R)- | 204.35 | 3.9 | 0 | 0 |
| 9 | 2-Butyl-5-methyl-3-(2-methylprop-2-enyl)cyclohexanone | 222.37 | 5 | 0 | 1 |
| 10 | Diethyl Phthalate | 222.24 | 2.5 | 0 | 4 |

For these selected phytochemical compounds, Lipinski’s rule of five parameters such as molecular weight, log P, and number of hydrogen bond donors and number of hydrogen bond acceptors were taken from the PubChem database (Table 3). From the compounds identified by GC-MS analysis, only those which obey Lipinski’s rule of five are alone subjected to docking experiment. Only compounds which fulfill Lipinski’s rule of five were used for molecular docking experiment. The structures of compounds from *Allophylus serratus* leaf extracts are shown in Fig. 2.

#### 3.2.3. Preparation of the protein

The target protein, which is cyclooxygenase-2 (Prostaglandin Synthase-2), three-dimensional (3D) structure was downloaded from the Protein Data Bank (PDB) database (www.rcsb.pdb) is give in Fig. 3a. The PDB database is a repository for the 3D structural data of large biological macromolecules such as proteins and nucleic acids. The PDB ID of COX-2 enzyme is 6COX-A, which is a complex of COX-2 enzyme with selective inhibitor compound SC-558. The active site region of the COX-2 enzyme where the inhibitor compound binds is given in Fig. 3b. This protein is a monotopic membrane protein which has two chains and 587 amino acids length. The ligands bound to this protein were HEM, NAG and S58 each having two chains.

**Figure 3:**
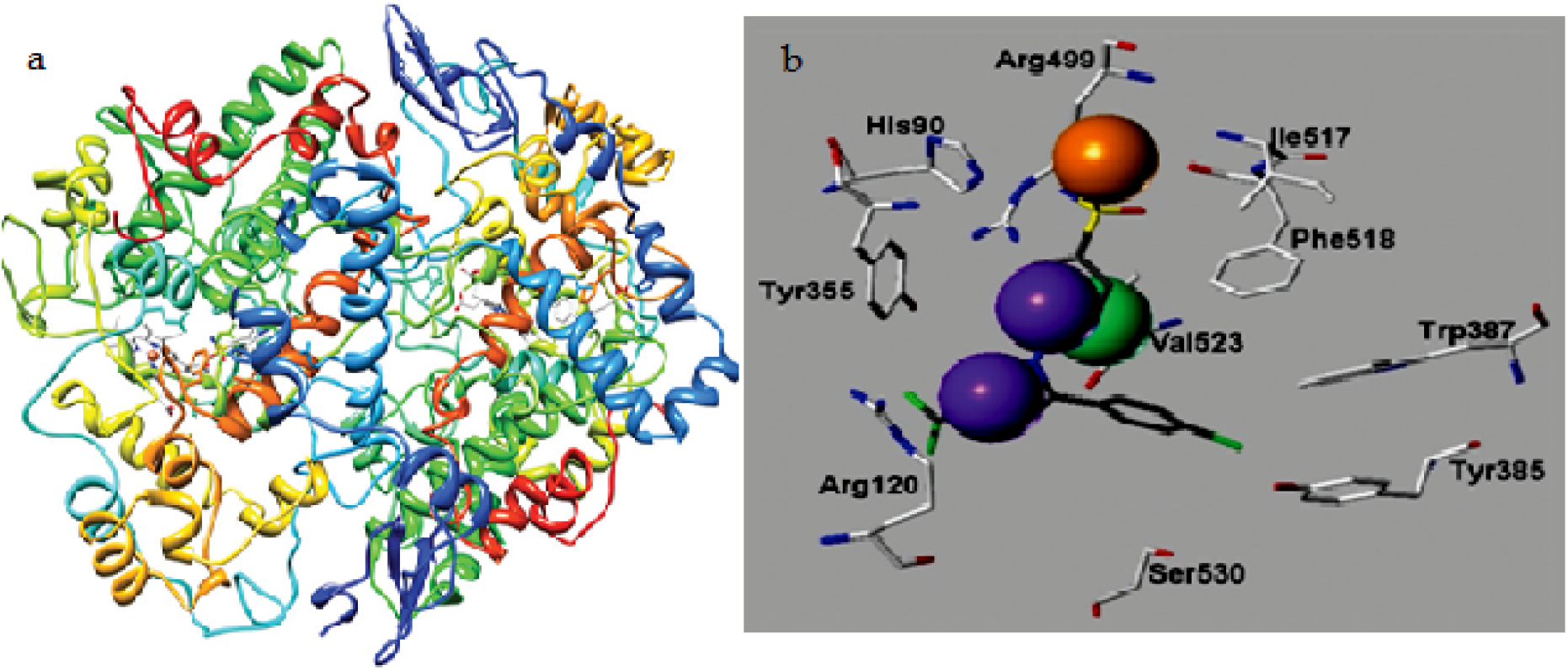
Cyclooxygenase-2 (COX-2) (a) Cyclooxygenase-2 (COX-2) (Prostaglandin Synthase-2) in complex with a COX-2 selective inhibitor SC-558 in I222 space group, (b) Cyclooxygenase-2 (COX-2) enzyme in complex withselective inhibitor SC-558 showing the active site region of COX-2enzyme

The protein 6COX_A Fig 7.4a was loaded into the Dockprep module of the UCSF Chimera software. Chain A of the protein was selected for preparation and docking. The active site of the protein is the binding pocket where reference ligand SC-558 is bound in this particular chain. Before docking, the protein crystal structures were cleaned by removing the hetero atoms such as, water molecules, unnecessary ligands and ions. Hydrogen atoms were added to correct ionization and tautomeric status of the amino acids and charges were added where required using Dock prep tool in the Chimera software. The docking process was performed using AutoDock Vina. Docking parameters were set following Autodock vina software procedures. The pdb coordinates of the protein and the ligand were submitted to AutoDock Vina. The binding energy and the binding contacts of each ligand were obtained. Analysis of the docked complexes was done using Discovery Studio 3.1 visualizer

#### 3.2.4. Docking

The biological activity of *Allophylus serratus* compounds against the COX-2 were evaluated using the 3D structure of the receptor retrieved from protein data bank site of COX-2 enzyme (PDB code: 6COX_A). For the selected compounds and protein the docked binding mode was established to link the docking score function.

The binding pattern analysis between COX-2 protein and ligands suggested that the binding pattern varied with the ligand nature. The docking results of the Declofenac and bioactive compounds from *Allophylus serratus* are given in Fig.4 and Table 4. The binding energy for the chosen phytochemical compound with the cyclooxygenase (COX-2) enzyme using Auto Dock vina is given in Table 7. 2. Docking studies show that the ligands bind to the active site region of COX-2 enzyme with good binding energy in the same hydrophobic pocket to that of Diclofenac control.

The docking results were represented in the form of e-negative values (Tables.2& 3). In the docking studies, higher negative e-values represent high binding affinity between the receptor and ligand molecules, indicating the higher efficiency of the bioactive compounds.

The results of the interactions of ligands (*Allophylusserratus* leaf extract compounds) with the Cyclooxygenase-2 (COX-2) receptor are summarized in Table 5.5. The docked ligands show scores ranging from −6.3 to −8.9. The docking score ofDiclofenac (control) against COX-2 was found to be-7.7 kcal/mol with two hydrogen bonds (Table 5).

**Table 5:** Molecular docking score for Secondary metabolites of *A. serratus*with COX-2

| SN | Ligand | Docking score (kcal/mol) | No. of hydrogen bonds formed |
| --- | --- | --- | --- |
| 1 | Isocaryophyllene | -6.5 | 0 |
| 2 | Sulfurous acid, dipentyl ester | -8.4 | 0 |
| 3 | Spiro[5.5]undeca-1,8-diene, 1,5,5,9-tetramethyl-, (R)- | -7.1 | 0 |
| 4 | Semicarbazide, 4-benzo[1,2,5]thiadiazol-4-yl-1-(5-nitro-furan-2-ylmethylene)- | -7.4 | 2 |
| 5 | 1,15-Pentadecanedioic acid | -7.2 | 2 |
| 6 | Illudol | -7.1 | 0 |
| 7 | 1,2-Benzenedicarboxylic acid, bis(2-methylpropyl) ester | -7.9 | 0 |
| 8 | 1H-Benzocycloheptene, 2,4a,5,6,7,8-hexahydro-3,5,5,9-tetramethyl-, (R)- | -8.9 | 0 |
| 9 | 2-Butyl-5-methyl-3-(2-methylprop-2-enyl)cyclohexanone | -7.7 | 1 |
| 10 | Diethyl Phthalate | -6.7 | 0 |
| 11 | Declofenac sodium | -7.7 | 2 |

Auto dock is a docking program used for virtual screening and predicting protein-ligand binding modes. It is an automated procedure for predicting the interactions of ligands with target protein. Table 7.5shows the molecular docking score of the secondary metabolites of leaf extract of *Allophylus serratus*. Out of the 44 compounds found in GC-MS analysis of methanolic and ethyl acetate leaf extracts, only 10 compounds that fulfill Lipinisks rule of five were docked with COX-2.

Out of the 10 components, 1H-Benzocycloheptene, 2,4a,5,6,7,8-hexahydro-3,5,5,9-tetramethyl-, (R)-has highest docking score(−8.9 kcal/mol) but with no hydrogen bonds. The result showed that other intermolecular interactions were predominantly involved in the interaction of the receptor protein and the ligand molecule. The second highest docking score with binding energy −8.4 kcal/mol without making any H-bond was obtained by Sulfurous acid, dipentyl ester with COX-2 followed by 1,2-Benzenedicarboxylic acid, bis(2-methylpropyl) ester which show the docking score binding energy of −7.7 kcal/mol. On theother hand 1, 15-Pentadecanedioic acid and the control Declofenac formed the maximum number of hydrogen bonds (2hydrogen bonds each) with the target protein. The least docking score binding energy of −6.3 was formed by Isocaryophillene with COX-2.

The docking models of the selected compounds(1) Isocaryophillene, (2) Sulfurous acid, dipentyl ester, (3) Spiro[5.5]undeca-1,8-diene, 1,5,5,9-tetramethyl-, (R)-, (4) Semicarbazide, 4-benzo[1,2,5]thiadiazol-4-yl-1-(5-nitro-furan-2-ylmethylene, (5)1,15-Pentadecanedioic acid, (6)Illudol, (7) 1,2-Benzenedicarboxylic acid, bis(2-methylpropyl) ester, (8) 1H-Benzocycloheptene, 2,4a,5,6,7,8-hexahydro-3,5,5,9-tetramethyl-, (R)-, (9) 2-Butyl-5-methyl-3- (2-methylprop-2-enyl)cyclohexanone and (10) Diethyl Phthalatein 3D view are shown in Figs. 7.5 and 7.6.

**Figure 7.5:**
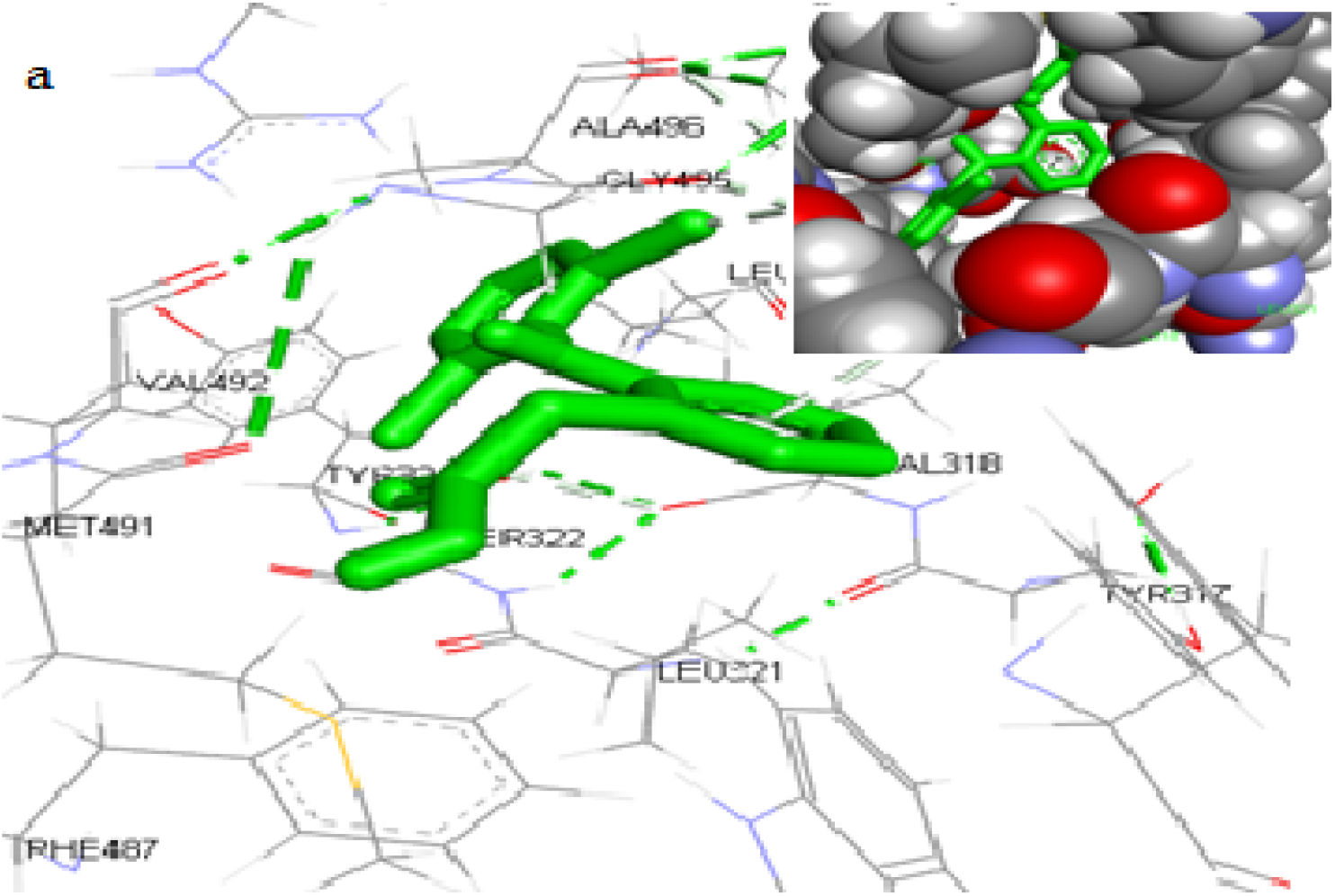
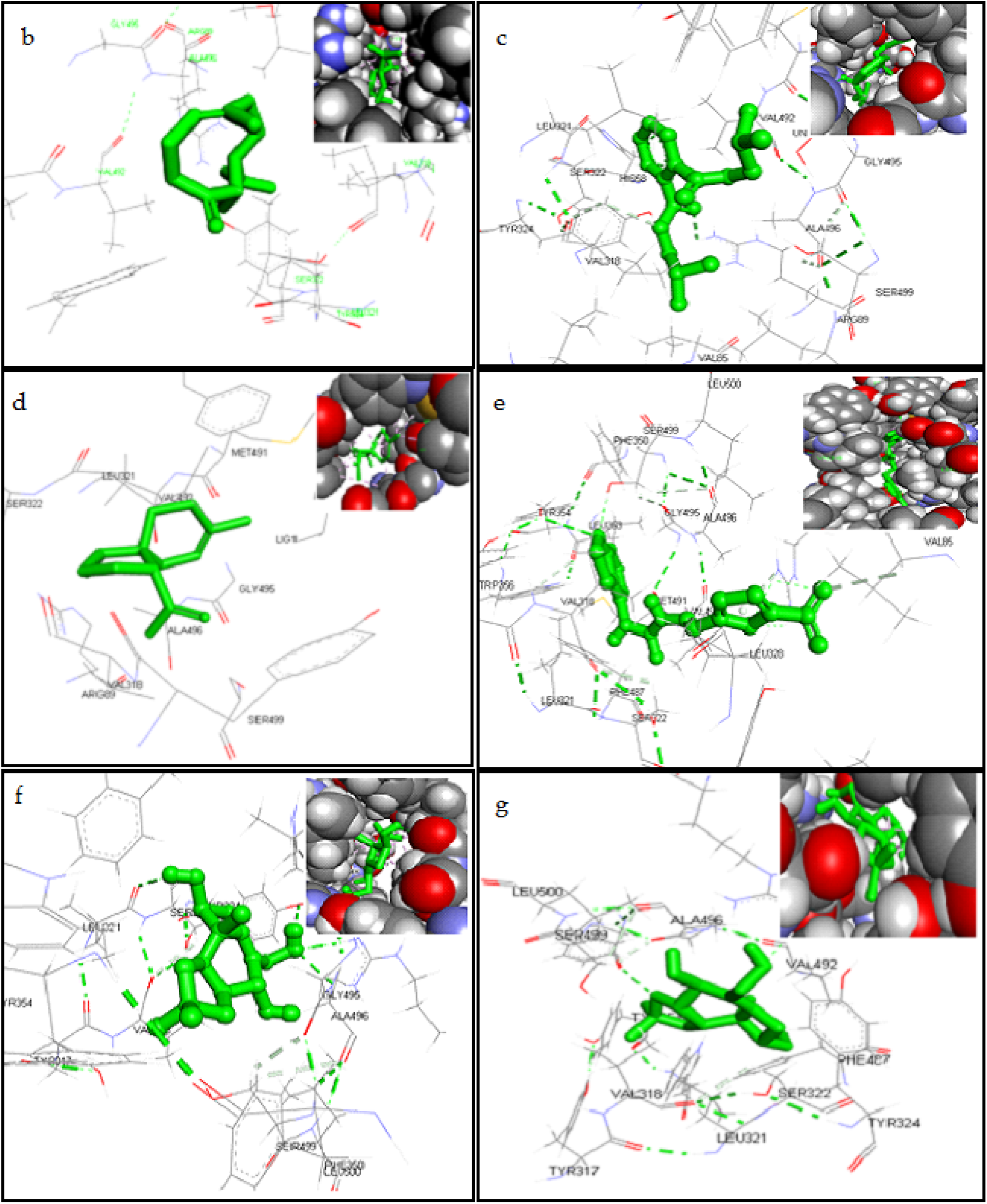
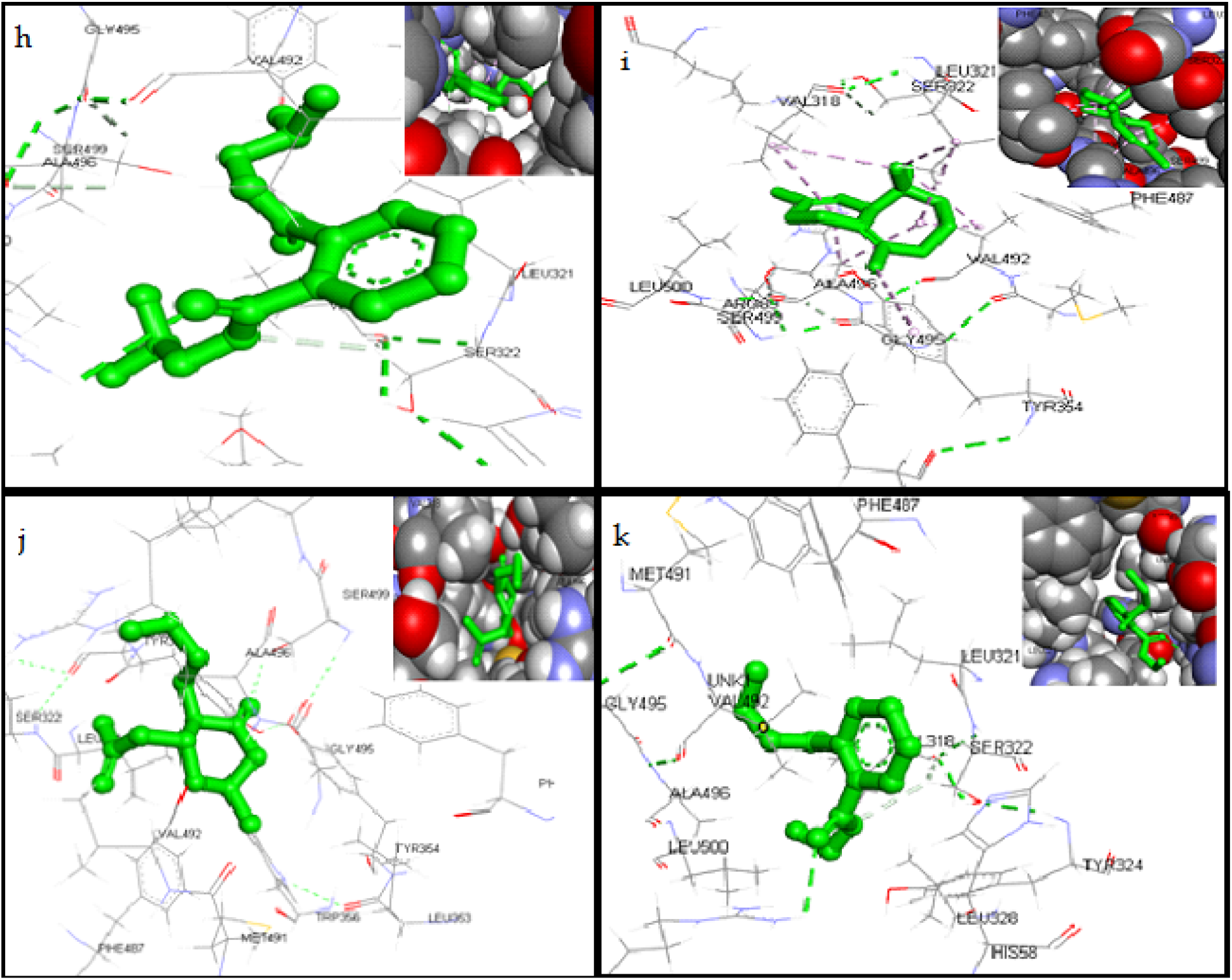
The Bioactive compounds from *A. serratus* leaf extract and Diclofenac docked with Cyclooxygenase (6COX-2) enzyme. a)Diclofenac, b)Isocaryophillene, c) Sulfurous acid, dipentyl ester, d) Spiro[5.5]undeca-1,8-diene, 1,5,5,9-tetramethyl-, (R)-,e) Semicarbazide, 4-benzo[1,2,5]thiadiazol-4-yl-1-(5-nitro-furan-2-ylmethylene)-, f) 1,15-Pentadecanedioic acid, g) Illudol, h) 1,2-Benzenedicarboxylic acid, bis(2-methylpropyl) ester i) 1H-Benzocycloheptene, 2,4a,5,6,7,8-hexahydro-3,5,5,9-tetramethyl-, (R)-, j) 2-Butyl-5-methyl-3-(2-methylprop-2-enyl)cyclohexanone and k) Diethyl Phthalate

**Figure 5:**
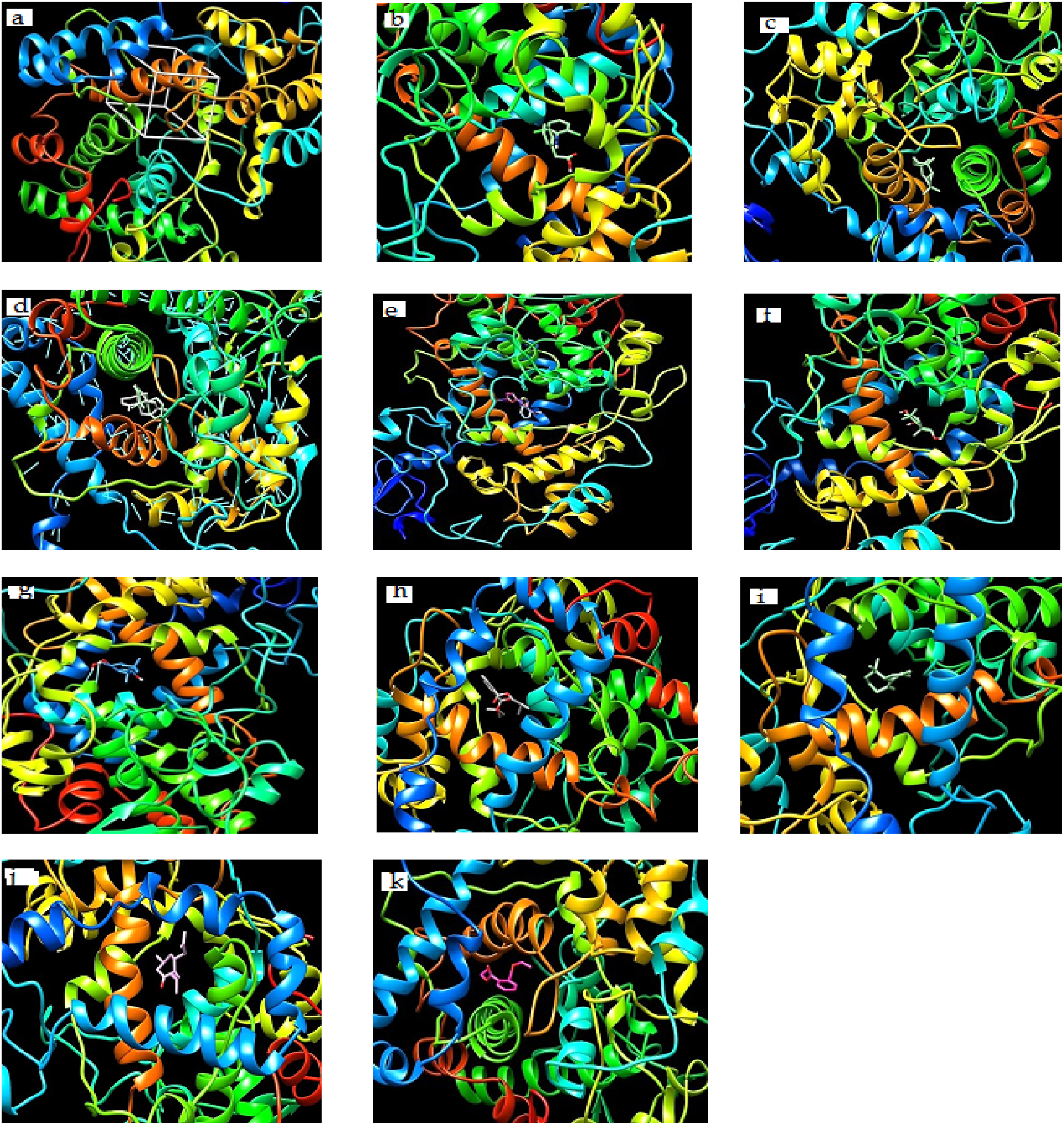
Binding modes of ten bioactive compounds from *Allophylus serratus*.(a) 6COX protein with active site (b).Isocaryophillene (c); Sulfurous acid, dipentyl ester (d); Spiro[5.5]undeca-1,8-diene, 1,5,5,9-tetramethyl-, (e); Semicarbazide, 4-benzo[1,2,5]thiadiazol-4-yl-1-(5-nitro-furan-2-ylmethylene, (f);1,15-Pentadecanedioic acid, (g)Illudol, (h) 1,2-Benzenedicarboxylic acid, bis(2-methylpropyl) ester, (i) 1H-Benzocycloheptene, 2,4a,5,6,7,8-hexahydro-3,5,5,9-tetramethyl-,(R)-(j)2-Butyl-5-methyl-3-(2-methylprop-2-enyl)cyclohexanone and (k) Diethyl Phthalate docked onto 6COX protein.

The results showed that all the bioactive compounds with target antigens produced high negative e-value. Thus, it is clear that the bioactive compounds were able to interact with any of the available binding sites of the COX-2 effectively. The above study clearly indicates that the bioactive compounds of *Allophylus serratus* were able to inhibit the activity of the COX-2 enzymes.

### 4. Discussion

The quantitative GC/MS phytochemical analysis of methanolic extract of *Allophylus serratus* leaf showed totally 32 compounds and that of ethyl acetate extract of *Allophylus serratus* leaf showed totally 12 compounds. These phytochemical compounds belong to various groups such as alkaloids, flavonoids, glycosides, saponins, tannins, phenol and terpenes. Flavonoids are phytochemical compounds which have antioxidant activity and are effective superoxide anions15scavengers. Alkaloids have antifungal properties because they have the ability to intercalate with DNA.

In order to understand the process of inflammation, it is important to understand the role of different chemical mediators that direct the inflammatory response (Wilson *et al*., 2006; Chopade and Mulla, 2010). Chemical mediators bind to specific receptors on target cells and can increase vascular permeability and neutrophil chemotaxis, stimulate smooth muscle contraction, have direct enzymatic activity, induce pain, or mediate oxidative damage (Wilson *et al*., 2006; Chopade and Mulla, 2010). Molecular docking is one of the most powerful techniques to discover novel ligands for proteins of known structure and thus plays a key role in structurebased drug design. The *in vitro* analysis of anti inflammatory activity of *Allophylus serratus* showed good results (Kero *et al*, 2017). The present study may act as supportive evidence that verify the anti-inflammatory properties of the *Allophylus serratus*, which may be because of the ability of phyto-constituents identified from this plant to inhibit various inflammatory mediators such as Cyclooxygenase-2.

Advances in computational techniques played important role in drug discovery process. To reduce the cost and time of drug discovery, virtual screening methods are routinely and extensively used. Virtual screening utilizes docking and scoring of eachcompound from a dataset and predicts the binding modes and affinities of ligand and receptor (Franca *et al*., 2013).Molecular docking has helped important proceedings to drug discovery for long time.Docking techniques help to recognize correct poses of ligands in the binding pocket of a protein and to predict the affinity between the ligand and the protein. At the end, docking describes a procedure by which two molecules fit together in three-dimensional space.

In this study, for the first time, phytochemical compounds from *Allophylus serratus* were docked with Cyclooxygenase-2 enzyme to see whether these compounds can bind to the active site of the enzyme and inhibit its activity so that confirm the anti inflammatory activity of the plant. The results of the docking study showed that all the tested phytochemical compounds (10 compounds) can bind to the receptor COX-2 similar to the control Diclofenac. This result clearly demonstrates that themolecular docking*approach*used was successful in finding novel COX-2 inhibitors from *Allophylus serratus* extracts.

In our present study, by means of AutoDock Vina, we docked 10 compounds from *Allophylus serratus* with active site ofCOX-2 enzyme and 1H-Benzocycloheptene, 2,4a,5,6,7,8-hexahydro-3,5,5,9-tetramethyl-, (R)-was found to provide mostsignificant binding score without hydrogen bond when compare to other compounds. Other compounds also showed significant binding score compared to the control Diclofenac. Moreover, the data obtained from docking is in agreement with previously reporteddata on synthetic compounds where amino acid residues associated with A chain of COX-2 protein wereinvolved for protein–ligand complementarily activity (Krishna *et al*., 2013).

